# The environmental context of the Middle-to-Late Stone Age Transition in eastern Africa: seasonality as a key factor

**DOI:** 10.1101/2024.12.09.627606

**Authors:** Marianna Fusco, Behailu Habte, Alice Leplongeon, Andrea Manica, Enza Elena Spinapolice, Michela Leonardi

## Abstract

In the transition between the Middle Stone Age (MSA) and the Late Stone Age (LSA) in eastern Africa, the archaeological record shows a gradual and asynchronous decline in MSA features and an increase in LSA characteristics. A link between this pattern and climatic variations has not yet been tested in the region using lithic attribute analysis.

To investigate that, we integrated technological data of blades and bladelets from eastern African contexts (Marine Isotope Stages 5–1) with large-scale paleoclimatic reconstructions. A principal component analysis (PCA) finds the first component (reflecting artifacts’ dimensions) significantly correlating with time. This highlights a progressive reduction in size over time, a trend that has already been suggested for the MSA-LSA transition. The second principal component reflects artifact shape and shows a significant correlation with the marked aridity of the dry season (a common proxy for seasonality in tropical regions), with higher specialization observed in more humid areas.

Based on this, we propose a new model where more variable blades reflect greater versatility in foraging strategies as adaptation to environments that become more challenging during part of the year. On the other hand, when it rains more during the dry season and differences through the year are milder, a more specialized toolkit with thinner, longer elements would emerge from refining and adapting to uniform and predictable situations and challenges.

## Introduction

Technological changes in lithic productions are often used to identify cultural shifts in prehistory (especially for the Paleolithic). However, the reasons behind technical and cultural transitions are highly debated, mostly because we know how tools change but we lack insight into the underlying reasons.

Such transitions do not always occur synchronously in different localities [1], and they may have a strong regional pattern, which complicates the definition of models for human groups’ behavior, adaptation, and evolution [2–4].

The transition from the Middle Stone Age (MSA) to the Late Stone Age (LSA) in Africa is an example of this problem that received considerable attention. It is defined on the basis of changes in material culture, particularly the appearance of artefacts traditionally interpreted as markers of increased cultural complexity (e.g. beads, bone tools, engraved objects); for a long time, the onset of the LSA in Africa has been framed as coinciding with the emergence of behavioral modernity.

This view was mostly based on what was suggested for Europe: the ‘revolutionary’ transition from the Middle to the Upper Paleolithic was linked to the distinction between *Homo sapiens* (i.e., “modern” humans) versus Neanderthals [5–11]. This generated a confusing superposition between the definition of the MSA-LSA transition and the beginning of “modern” human behavior [5]. However, current evidence shows that the boundary between the MSA and the LSA is not abrupt, and it is now widely accepted that this transition is more of a gradual process than a sudden “revolution” marking the shift to a more “complex” behavior [3,12]. Now, the link between behavioral complexity and LSA-makers is considered obsolete, and the archaeological evidence shows an asynchronous presence and duration of innovations across the African continent [12]. In fact, evidence of what is defined as “behavioral complexity” is present in Africa as far back as >100 ka [13–15], in a fully “MSA environment”, and even earlier [16]. Moreover, evidence for great regional variation in late MSA and early LSA assemblages suggests that there were multiple local or regional transitions, each with their specific expression[17]. In eastern Africa, such processes happened in the Late Pleistocene, between ∼70 and 15 ka, i.e. Marine Isotope Stages (MIS) 4-2[2,4,18–20].

The MSA, by definition, is characterized by the presence of the Levallois and discoid methods, as well as retouched and unretouched points (considered the hallmarks of formal MSA tools) [21]. Its chronological range spans from ∼300 to ∼30 ka [3,12,16]. The emergence of the MSA can be considered contemporaneous to the emergence of the first *Homo sapiens* [22,23], but some of the traits considered typical of the MSA are occasionally found in Holocene contexts e.g. the presence of the Levallois method for flake production[19,24–27]. In contrast, the LSA is characterized by - depending on the regions - all or some of the following features: (a) increased production of blades and bladelets using volumetric reduction methods, (b) more frequent use of the bipolar technique on anvil (e.g. for the production of unretouched bladelets, or quartz flakes/bladelets to be transformed into microliths), (c) increase in the production of microliths and backed tools, (d) more frequent use of ochre and other ornamental objects (e.g. shell beads), and (e) preference for fine-grained raw materials [2,4,20,28–32].

The asynchronous emergence of the LSA across the African continent offers a great opportunity to investigate the drivers behind these changes, but until now several challenges have prevented a unified view of their mode and tempo [2,12,17].

First, the general models of Late Pleistocene archaeology in eastern Africa have been elaborated mainly through the analysis of single contexts, making it extremely difficult to develop a broader contextualization of the changes observed between MIS 4 and 2.

Moreover, research efforts are strongly biased toward specific parts of the continent, both because of long-standing research traditions and practical challenges (e.g. political instabilities).

Also, the lack of long-stratified and well-dated sites providing a comprehensive view of the events between ∼80 and ∼17 ka makes it extremely difficult to explore the origin and nature of the transition at a broader scale. For example, in Panga ya Saidi, Kenya, between the layers attributed to MSA and LSA there is an increase in small-sized lithic productions. However, the selection of finer-grained raw materials in the LSA layers are here specifically considered as the hallmark of the LSA [20,31,33,34]. In Nasera, Tanzania, demographic change may have been linked to a decreased residential mobility across the MSA/LSA transition, in the context of environmental fluctuations, but with no evidence for a particular selection of different raw materials from the MSA to the LSA[32]. However, in these multi-layered archaeological contexts, such patterns do not display a regional tendency. Such trends have only been recognized when compared to the MSA levels in single-layer contexts. This has led to an incomparability of the data, challenging the validity of explanatory models at a large scale for the MSA-LSA transition.

The analysis of material culture details human choices, but more is needed to identify the drivers behind them. To do so, it is necessary to look at large-scale human behavioral dynamics, performing comparative analyses on lithic assemblages from wider regions and integrating different data (i.e. environmental, genetic, climatic)[35]. One possible explanation to the observed pattern in the MSA-LSA transition is that the emergence of LSA toolkit (e.g. microliths and backed pieces) could be a response to changes in the environmental conditions during the Late Pleistocene [2,4,30,32]. Such hypotheses can only be tested integrating archaeological and paleoclimatic data[36].

Environmental conditions, including temperature, the amount of rainfall, and the differential availability of natural resources, are likely to have played an important role in shaping how past societies behaved and adapted through time. For example, refugia, physical barriers, and corridors led to population bottlenecks and expansions and have driven the emergence, development, and spread of past human behaviors and cultures[37–40].

Eastern Africa is particularly suitable for integrating archaeological and paleoclimatic data, as it includes various habitats that might explain distinct regional traditions in lithic productions [40]. In eastern Africa the climate is mostly arid between MIS 4 and MIS 1 but includes significant oscillations and humid episodes within MIS 4 and 3[41]. MIS 2 is marked by the so-called “Big Dry”, occurring around ∼23-20 ka and roughly corresponding to the Last Glacial Maximum (LGM). Humid conditions are observed both before and after the LGM, i.e., before ∼24 ka and after ∼15 ka[42]. During MIS 1 we have evidence of the African Humid Period (AHP), characterized by increased precipitation in eastern Africa and improved environmental conditions, resulting in a more humid environment[42,43].

At a local level, the sub-region of the Horn of Africa is characterized by aridity during MIS 4, with wetter intervals. During MIS 3, a more humid phase occurred, especially ∼50-40 ka, with some arid intervals. MIS 2 in the Horn is characterized by episodes of aridity, with peaks (not only corresponding to the LGM) in which precipitation decreased to up to 25% [42–45]. The region also experienced the impacts of Heinrich Event 1 (around 16 ka), a period of significant cooling and aridification due to massive icebergs breaking off from glaciers in the North Atlantic [41,46]. This event further intensified arid conditions in eastern Africa, contributing to the drying up of key water bodies like Lake Victoria and Lake Tana. Finally, the beginning of the AHP, which is a general wet phase, in the Horn was delayed by the Heinrich event, which prolonged the aridity until around 12 ka[41,42].

Different technological changes in lithic production are observed during these fluctuations. Among them, a gradual increase of different technical traditions (especially between the end of MIS 3 and the onset of the MIS 2) in the Horn of Africa [2,17], while in Kenya this phenomenon is placed earlier in time [20,31]. As technical changes happen variably in space and time, it is not clear whether the transition from the MSA to the LSA reflects cultural and behavioral adaptations to large-scale climatic and environmental changes, or responses to localized events that spread to larger areas.

Integrating different data (i.e., lithic attributes and paleoclimate) has already given interesting results, showing a complex scenario for human occupation and dispersal dynamics in North Africa during the MIS 5[47]. Similar approaches have been proposed for eastern African contexts, but lacking information about the technological process of lithic artifacts production [48]. It is most likely that the MSA-LSA transition was linked to a variety of factors (e.g. large-and small-scale climatic fluctuations, changes in demographic patterns).

In this article we test if and how paleoclimate shows a significant correlation with the technological changes observed during the final MSA and the onset of LSA in eastern Africa. This paper does not intend to provide a direct explanation for how different blade shapes affected hunting strategies in variable environments, nor does it aim to establish causality between climate and toolkit morphology. Rather, we present this as a first step toward identifying broader patterns of association.

## Results

### Data analyzed

We considered the lithic assemblages from each dated layer of five archaeological sites dated between MIS 5 and MIS 1 in Ethiopia – Goda Buticha (GB), Porc Épic (PE), Gotera 10 (GOT-10), Deka Wede 4 (DW4) – and Southern Kenya – Enkapune ya Muto (EYM)[19,28,49–52] (Figure 1, table 1 and Materials). We focused the analyses on blades and bladelets (flakes in which the length is at least twice the width, also defined as “elongated blanks”). During the MSA laminar reduction strategies are aimed at producing both blades and bladelets, with an occasional presence of backed pieces [17]. In the LSA, their presence increases, as most of LSA assemblages are characterized by a marked emphasis on their production for the manufacture of formal tool types such as backed pieces and geometrics [51,53]. For this reason, elongated blanks represent a particularly suitable analytical category for evaluating technological variation across the MSA–LSA transition[4,54,55]. In addition, elongated blanks are the lithic category for which the most data were available in the dataset, allowing us to perform a more comprehensive analysis.

**Figure 1:**
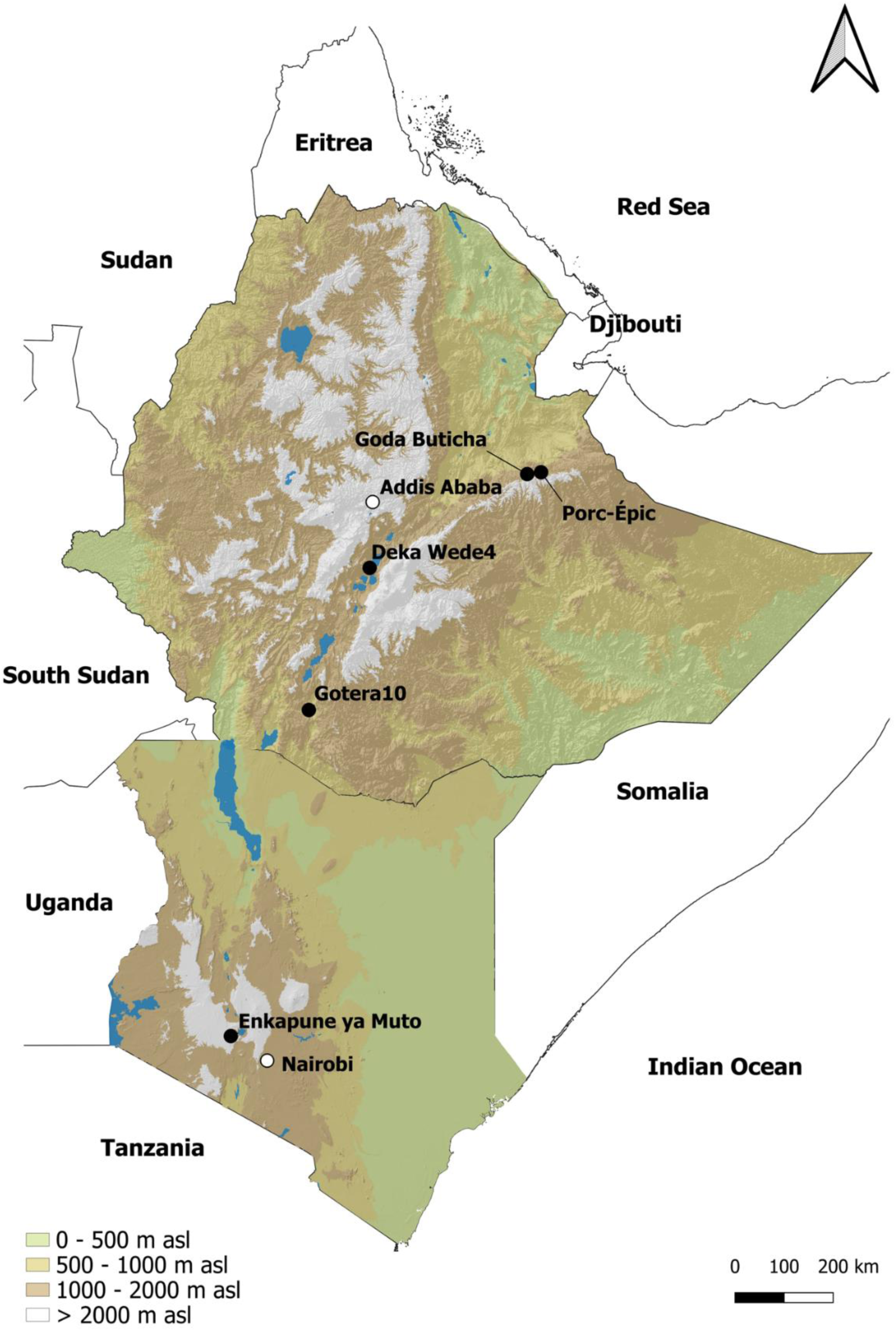
map showing archaeological context considered. 1: Porc Epic; 2: Goda Buticha; 3: Deka Wede4; 4: Gotera; 5: Enkapune ya Muto. Map created with QGIS 3.34.11.

**Table 1:** list of sites considered in the analysis with the identification of chronology, cultural attribution and number of elongated blanks sampled. More information about the chronology (including the original references) are found in the Methods section.

| **Site** | **Layer** | **Chronology** | **Cultural Phase** | **Sample size** | **Number in Figure 2** |
| --- | --- | --- | --- | --- | --- |
| **Goda Buticha** | IIIdIf (030-000) | 63000±7000 years ago | MSA | 14 | 4 |
| **Goda Buticha** | IIIdIf (100-030) | Between 29680+/-230 and 42500+/- 1000 Cal BP | MSA | 131 | 5 |
| **Goda Buticha** | IIc (160-100) | Between 5590+/- 50 and 6940+/-40 Cal BP | Middle Holocene | 186 | 10 |
| **Goda Buticha** | I (190-160) | 3725+/-30 Cal BP | Late Holocene LSA | 35 | 11 |
| **Porc Epic** | IIIId-IIIc | Between 130000 and 60000 years ago | MSA | 92 | 1 |
| **Porc Epic** | IIIb-IIIa | Between 130000 and 60000 years ago | MSA | 167 | 3 |
| **Porc Epic** | II | Between 130000 and 60000 years ago | MSA | 29 | 2 |
| **Gotera 10** | SU6 - SU7 | Between 45760±540 and 43270±390 Cal BP | LMSA | 48 | 6 |
| **Deka Wede 4** | 1500 | Between 26910± 100 and 41640±420 Cal BP | LMSA | 102 | 7 |
| **Enkapune ya Muto** | GG1.2-3-4-5-7 | Between 55000 and 45000 years ago | LSA | 46 | 8 |
| **Enkapune ya Muto** | DBL1 | 47166-41781 | LSA | 126 | 9 |

### Morphological trends

For each elongated blank we collected lithic attributes (e.g. metrics about their shape and size) suitable to compare different collections [56] (see Methods and table 1).

We first performed a Principal Component Analysis (PCA) to evaluate the differences between the MSA and the LSA assemblages (Figure 2: A). In the PCA, the two first principal components accounted for 68.3% of the overall variance. PC1 is mostly influenced by dimension maximum length (XTL), maximum width (XTB), and maximum thickness (XT). On the other hand, PC2 is driven by the elongation index (EI) and flattening index (FI). Following this, variation along PC1 reflects changes in size, with smaller blades and bladelets on the right of the plot, and larger ones on the left. Similarly, the value of PC2 captures the shape, with longer and thinner blanks on bottom and shorter and thicker ones on the top of the PCA (Figure 2).

**Figure 2:**
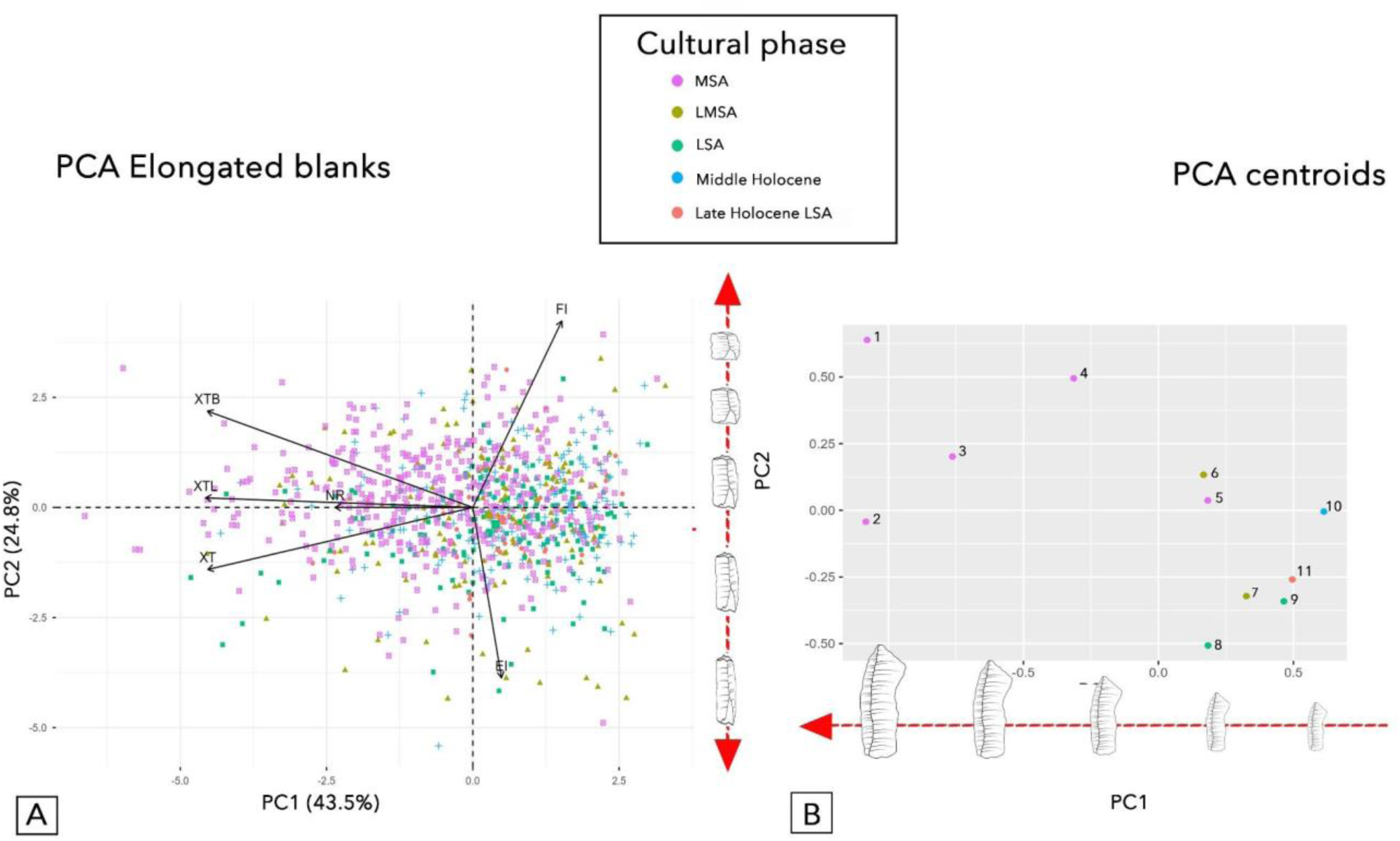
A: biplot with PCA results grouped by cultural phases and showing both individuals and variables; B: PCA showing centroids grouped by cultural phase. Cultural phases: Middle Stone Age (MSA), Late Middle Stone Age (LMSA), Late Stone Age (LSA), Middle Holocene and Late Holocene LSA. Numbers indicate each individual archaeological layers. MSA: 1: Porc Epic IIId-IIIc; 2: Porc Epic II; 3: Porc Epic IIIb-IIIa; 4: Goda Buticha IIdIIf (63ka); 5: Goda Buticha IIdIIf (44-31 ka). LMSA: 6: Gotera 10; 7: Deka Wede 4. LSA: 8: Enkapune ya Muto GG1; 9: Enkapune ya Muto DBL1. Middle Holocene: 10: Goda Buticha. Late Holocene LSA: 11: Goda Buticha.

To better visualize the distribution of lithic assemblages within the PCA space, we summarized the information for each archaeological layer by calculating their centroids: i.e. the average position of all their artifacts (Figure 2: B, Methods). The centroids show a progressive size reduction through time (Figure 2: B). The MSA assemblages (Porc Épic IIIa-d and II layers, Goda Buticha IIdIIf 030-000 (63ka) and IIdIIf 100-030 (∼44-31ka) layers, (Figure 1: B nr 1-5) have the largest size. In contrast, Gotera (SU6 and 7, ∼45ka) and Deka Wede4 (SU1500, ∼30ka) appear on the left side of the graph with smaller artifacts (Figure 1: B, nr 6 and 7). The GG1 layer of Enkapune ya Muto (Figure 2: B, nr 8), dated to ∼50 ka tends to be smaller than the Late Middle Stone Age (LMSA) artifacts from Gotera, but bigger that Deka Wede4, even if it is defined as LSA. Enkapune ya Muto DBL lithics (∼39ka) (Figure 1: B, nr 9) are smaller than MSA and LMSA ones (as well as GG1 LSA assemblage), but bigger than the Holocene data from Goda Buticha (layers IIc ∼8ka and I ∼2-0.7ka) (nr. 10 and 11 in figure 2: B). It must be noted that the superimposition of these data in the PCA reflects the blurry boundaries between the LMSA and the beginning of the LSA in the region in terms of blade/bladelet dimensions. It is also worth highlighting the importance of variability between different research traditions and classification of lithic assemblages: the difficulty in distinguishing LMSA from early LSA may also be due to this. This supports the growing consensus that the transition from the MSA to the LSA was not abrupt or reflected a “revolution”, but rather a gradual process characterized by technological and behavioral continuity alongside innovation [3,4,12,57].

Despite the classification of MSA and LSA for the assemblages considered in this study, the Enkapune ya Muto level GG dated to ∼50ka and defined as LSA, in the PCA (Figure 2: B, nr 8) appear to be an outlier compared to the other more recent LSA cultural phases and the level DBL1 of the same site. However, when comparing the technological changes with the chronology, its position in the plot seems consistent, beyond its cultural attribution (as confirmed by the correlation between PC1 and time).

The PC1 in our results clearly distinguishes between the MSA, LMSA, LSA and Middle-Late Holocene data showing a progressive reduction in size from ∼130 ka to the Holocene in the sites analyzed.

The distribution along PC2 shows a different pattern. In Figure 2: A, both the MSA and LMSA assemblages (purple and olive-green dots, respectively) exhibit a very large morphological variability. Among them, LMSA tends to have more elongated and thinner blades and bladelets, while the MSA has shorter and thicker ones. On the other hand, in figure 2: A, LSA assemblages (in green, while Holocene assemblages in blue and red) appear much more homogeneous, with a tendency to have similarly thin and elongated blanks, with LSA at the lower corner of the graph. This reflects another trend beyond a simple reduction in sizes [2,19].

The middle and late Holocene LSA Goda Buticha IIc layer (Figure 2: B, nr 10 and 11) represent an exception to the trend described above, probably due to the co-occurrence of both MSA and LSA characteristics [19]. These results prompted to investigate in more depth the reason behind these patterns.

### Raw material selection as potential driver of changes

Raw material selection is among the most common drivers behind technological changes [20,33,58–60]. Therefore, we tested if different raw materials affected the shape or size of the analyzed assemblages.

There is evidence of several sources for the materials utilized within our sites (see Materials and Methods).

Similarly to what we did above, we calculated for each assemblage the centroids for the different raw materials over the first two principal components (Figure 3: A).

**Figure 3:**
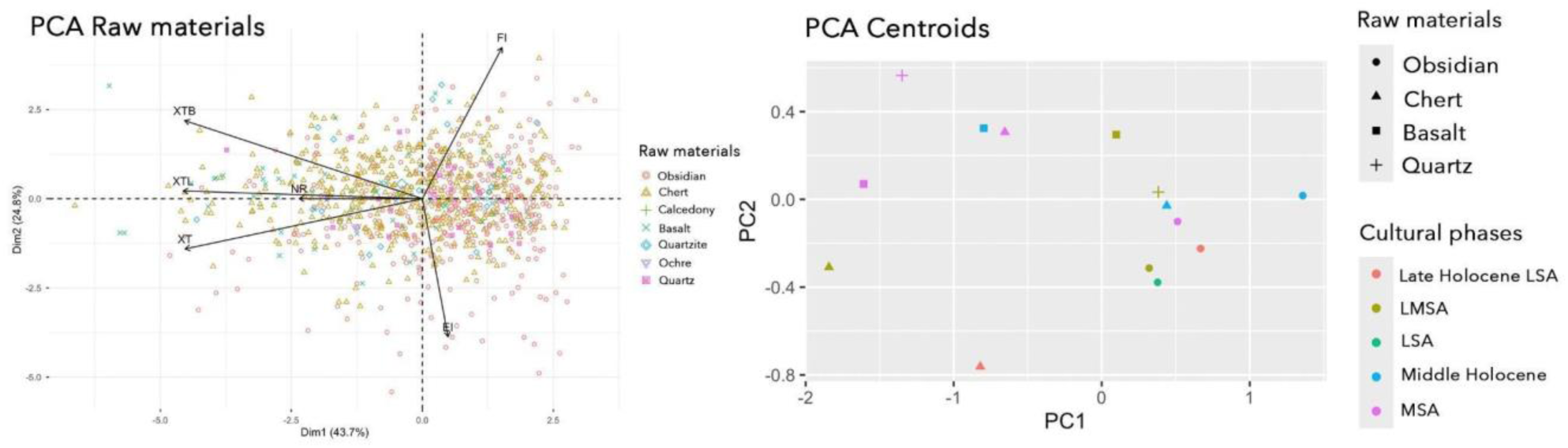
PCA showing each elongated blanks and variables subdivided by raw materials. Variables used for the PCA are elongation index (EI), flattening index (FI), Maximum breadth (XTB), maximum length (XTL), maximum thickness (XT) and number of dorsal scars (NR). (A). Plot showing the distribution of PC raw material centroids indicating also cultural phases (B).

The average artifact sizes of obsidian, chert, and quartz artifacts follow the expectations for their chronology and cultural phases: larger during the MSA and show a progressive reduction in later phases (Figure 3: B).

However, raw materials such as basalt do not display a consistent pattern of size reduction in relation to chronology: some large basalt artifacts are attributed to the Holocene or other cultural phases (Figure 3: B). This homogenous use of the different raw materials over time supports the idea that it is not the choice of materials the driver of the technological trends observed in our data,

### Paleoclimate as potential driver of changes

We then tested the hypothesis of paleoclimate as a potential driver for the technological changes observed in our data. We linked each assemblage to paleoclimatic variables reconstructed at a large scale[58] for its location and age using the *pastclim* package[59] in R (see Methods for additional details). Reconstructions are available at millennial/bimillenial intervals with a spatial resolution of 0.5° × 0.5°, from which the climatic variables for the analyses are directly derived. Through this approach, we can obtain consistent and comparable climatic estimates for all contexts analyzed, irrespective of the level of conservation of other proxies. Given the wide chronological and geographic extent of this study, we believe that such a resolution is adequate to recognize large-scale trends and patterns [61,62]

We tested annual measures of temperature (annual mean) and precipitation (total annual), net primary productivity, and the precipitation of the driest season, which is the most likely limiting factor associated with high seasonal variations in tropical environments. The seasonality of resource acquisition has been recognized as a key aspect of hunter-gatherer subsistence behavior as well as social and economic organization[60]. It is well known that the adaptation to different seasonal cycles observed in modern hunter-gatherer populations has shown that a wide range of factors related to technology, culture, and subsistence strategies shift in response to seasonal environmental changes[63]. However, in eastern Africa, few studies[64] have explicitly examined how the climate varies through the year as a potential driver of technological change among Pleistocene hunter-gatherers. For this reason, we chose to test this specific variable in our analysis. Because of the large chronological uncertainty of some of the layers, we did not use mean/median ages. Instead, for the whole list of layers, we resampled 1000 times a set of dates from each relevant range [65]. For each of these 1000 sets of resampled dates, we calculated Pearson’s correlation between the positions of the centroids of each layer on PC1 and PC2 and five variables. These include the chronology (time) and four measures summarizing the overall environmental characteristics: mean annual temperature and total annual precipitation to take into account the overall fluctuations in temperature and rainfall, net primary productivity to represent the type of environment, and the precipitation during the driest season as the ecologically limiting factor linked to differences in seasonality (Figure 4). As an aggregated measure of correlation, we used the mean ρ (rho) over the 1000 chronologically resampled sets, and we calculated the significance as the number of them in which the p-value was below 0.05 [65].

**Figure 4:**
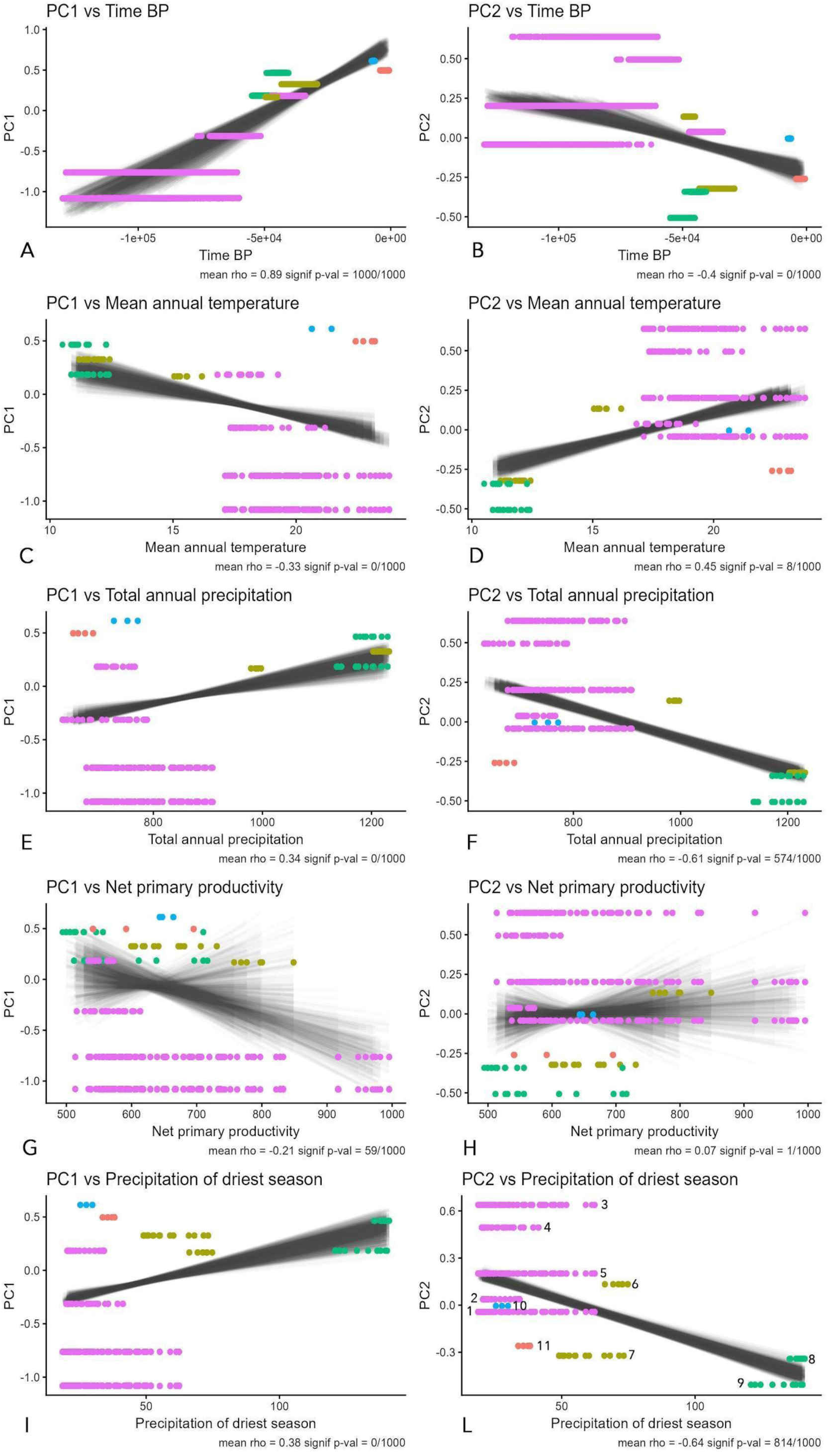
Integration of paleoclimate data to test the role of climate and environment within the MSA/LSA transition. The first column shows the correlation between PC1 and the different variables tested, while the second column focuses on PC2. Each line represents a different variable, from top to bottom: time, mean annual temperature, total annual precipitation, net primary productivity and precipitation of the driest season. Numbers in boxes L indicate each individual archaeological layer. MSA: 1: Porc Epic IIId-IIIc; 2: Porc Epic II; 3: Porc Epic IIIb-IIIa; 4: Goda Buticha IIdIIf (63ka); 5: Goda Buticha IIdIIf (44-31 ka). LMSA: 6: Gotera 10; 7: Deka Wede 4. LSA: 8: Enkapune ya Muto GG1; 9: Enkapune ya Muto DBL1. Middle Holocene: 10: Goda Buticha. Late Holocene LSA: 11: Goda Buticha.

As already suggested by Figure 2: B, PC1 (reflecting the size of the blanks) is significantly correlated with time (Figure 4: A), even after the chronological resampling (mean rho=0.89, all resampled sets significant). PC2, reflecting the shape uniformity of the lithics analysed, does not show significant correlation with mean annual temperature (rho=0.45, 8/1000 repetitions significant) and net primary productivity (rho=0.07, only 1 repetition significant). Total annual precipitation, instead, shows a higher correlation (r=-0.61), with more than half of the repetitions significant (574/1000). When looking at the seasonal aspect of rainfall (precipitation of the driest season), the correlation is higher (rho=-0.64) and overall significant (814/1000 repetitions) (Figure 4: L).

Elements with a higher flattening index (i.e. with higher values of PC2) are found in locations with larger differences between seasons, while elements with a higher elongation index (lower values of PC2) are associated with a climate that is more uniform over the year.

To check for the potential influence of time on our results, we tested for its correlation with all variables considered, finding low values of rho (figure 5). The correlation between the two measures of precipitation is expected and reflected in our results (total annual precipitation and precipitation of the driest season). Similarly expected is the high correlation between both these and mean annual temperature: fluctuations in rainfall through such long periods are linked to glacial cycles similarly to what we observe for the overall temperature. It must be noted that the mean annual temperature does not appear to correlate significantly with PC2, suggesting precipitation (more specifically its seasonality) as the main variable mirroring the characteristics in our data.

**Figure 5:**
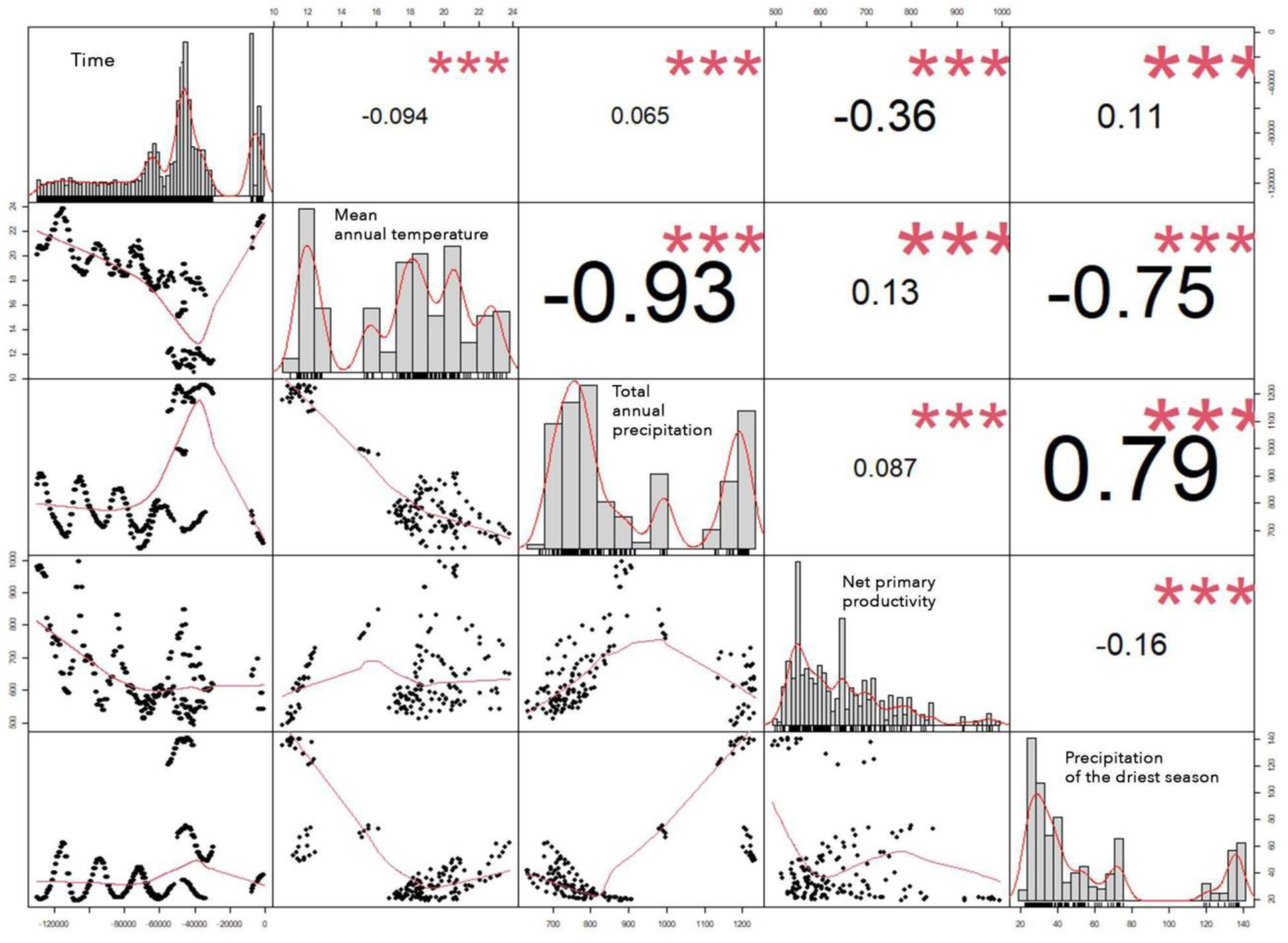
plot showing the cross-correlation between all variables analysed: time, mean annual temperature (bio01), total annual precipitation (bio12), net primary productivity (npp), precipitation of the driest season (bio17).

## Discussion

In this article, we perform computational analyses on elongated blanks from eastern African contexts dated to Marine Isotope Stages 5 to 1 to identify chronological trends.

After identifying in our dataset the principal components representing the overall size and shape homogeneity of our samples, we tested for correlation between these two measures and different variables reflecting chronology and environmental conditions. Our results suggest a relation between seasonal variation and lithic attributes, which is consistent with the observation that modern hunter-gatherer groups often adjust their cultural and technological behaviors in response to wet and dry seasons (see below).

It is important to highlight that such correlations do not imply causation. At the same time this kind of analysis can be valuable in identifying specific patterns to be tested empirically, which is the approach that we follow in this work. This is the reason why our results are presented here as a theoretical framework that requires further testing.

Variations in sample size may influence the results presented here; therefore, we conducted several tests to examine this possibility (see section on “*Check for an effect of sample sizes over the results*”).

We first examined whether smaller sample sizes were associated with reduced diversity in blank size and shape. If such a relationship existed, it would suggest that limited samples might only capture a narrow portion of the overall variability. However, our analysis does not support this hypothesis.

As a second step, we investigated whether sample size or technological diversity correlates with the resampled ages of the assemblages. A significant association would have implied that the observed patterns in variability could have been influenced by temporal trends or time-dependent preservation biases. Yet, these hypotheses are also not supported by the data.

Our results show a clear trend toward the reduction in the size of elongated blanks through time; a pattern more evident for assemblages classified as MSA (Figure 2: B). However, if we consider the cultural label classification, the reduction in size started before the MSA/LSA transition and it follows a chronological order that goes beyond the cultural definition. For example, the Early LSA GG1 layer from Enkapune ya Muto (8 in figure 2: B), has on average bigger blades and bladelets than the LMSA assemblage of Deka Wede 4 and similar values to the LMSA of Gotera 10, both more recent.

While a reduction in size of blanks, cores, and tools in the late Pleistocene has been commonly reported, the reasons behind such choices are highly debated [17,35,71]. As already discussed, size reduction was one of the characteristics that defined the LSA and differentiated it from the MSA[20,29,34,66,67], but it is now accepted that small size alone is not enough to categorize an assemblage as LSA [17].

Small-sized artifacts (<30-50 mm) are documented in various contexts, even older than LSAs [66–68]. In eastern Africa, they are found at Omo Kibish ∼200ka, Herto dated ∼150ka and Aduma, ∼80ka, as well as at Panga ya Saidi ∼67ka [20,31,34,69,70]. In South Africa, evidence of small flakes and blades is present as early as in the MSA, sometimes associated with the Howiesons Poort industry (characterized by small geometric tools used probably as inserts in multi-component tools) [71–73].

Our results show a clear trend of progressive size reduction in both blades and bladelets through time within the contexts considered in this paper, spanning from the MSA to the Holocene LSA. On the other hand, the incoherency between the definition of MSA-LSA transition and such size changes should push us to consider this phenomenon as something not marking the transition itself, but more of a chronological trend.

Various hypotheses have been proposed regarding the shifts in artifact sizes, accompanied by ethnographic analyses in contexts from both North America and Australia [66,74], including symbolic aspects related to the miniaturization of flakes and blanks in general [34,74]. Additionally, they consider factors associated with raw material economics in the contexts of population growth, reduced residential mobility, and the cost-benefit strategy of producing a greater number of smaller tools to maximize efficiency [66,68,74]. In many contexts, raw material selection played a significant role in lithic production [75–77]. Pargeter and Shea [66] suggest that some raw materials, (e.g. quartz) are easier to work into smaller-sized artifacts. There is evidence of technological changes in eastern African industries during the Late Pleistocene and early Holocene being associated with the exploitation of different raw materials[20,31]. The transition from the MSA to the LSA has been suggested to be linked to the selection of finer-grain raw materials, better suited for small tools[20,33]. While this evidence has been tested on single sites [20], our results do not support it on a larger spatiotemporal scale.

As mentioned, extensive discussions have occurred on the ecological and environmental aspects of lithic miniaturization and, in general, technological changes in stone tool use and production [28,68,71,78]. In eastern Africa during the Late Pleistocene, a series of environmental changes led to oscillations between arid and humid periods [41]. Based on our results, we propose that the correlation between the shape of blades and bladelets (i.e. elongation and flattening) and seasonality may suggest adaptive strategies in response to different degrees of resource predictability (e.g. raw material availability, seasonality of the game, water availability).

In this model, unpredictable environments are suggested to be linked to reduced resource reliability rather than climatic irregularity. Severe dry seasons, even when seasonally predictable, can amplify ecological uncertainty by constraining and spatially redistributing critical resources (e.g. water, fauna). We therefore refer to situations where the timing, location, and abundance of key resources fluctuate in ways that cannot be reliably anticipated by human groups, requiring flexible technological and mobility strategies to mitigate the risk of resource failure [79,80].

This has already been proposed as a driver for stone tool production and function since older periods [81,82].

Human response to climatic changes and environmental conditions during the Pleistocene is reflected in their ability to adapt and live in different ecologies while habitats expand or shrink. The diversity in lithic production and the different outcomes of technical evolution within these groups likely result from a modification and/or adaptation of hunting, butchering, animal processing strategies, and other subsistence practices based on the surrounding environment. It is widely accepted that resource predictability (a consequence of more stable environments) has impacted hunter-gatherer behavioral dynamics, in terms of subsistence, settlement, and mobility strategies[60,83–86]. For example, the exploitation of raw materials to anticipate inadequate conditions is proposed as one of the features that differentiates the concept of curation (intentional maintenance, reuse, and prolonged use-life of stone objects) from the one of expediency (minimal technological effort in object production)[85]. Going beyond the multiple definitions of these concepts [87–92], it was proposed that curated objects are produced where the need to cope with unpredictable environmental conditions leads to an anticipated preparation of raw materials. Objects were curated because there was no guarantee of always having the necessary raw materials available, in this way time was invested in advance [93–95]. Conversely, in favorable conditions, this anticipation may have been not necessary, with no time invested in advance, with expediency reflecting highly predictable resource availability [60,91,95].

Similarly, for our results, we hypothesize that in periods in which the dry season was less harsh, the prevalence of longer and thinner elongated blanks could be linked to predictable resource availability: a specialized toolkit. In regions with relatively stable environmental conditions, we expect that animal (and vegetable) resources would be available for the whole year. In such a scenario, hunter-gatherer groups would have the opportunity to rely on a stable toolkit for activities such as hunting and fishing, which would require homogeneous, longer and thinner blades [85–87,89,96].

On the other hand, a preference for more variable, shorter, and thicker blades may be a response to an environment where the dry season is more challenging and resource availability is relatively unpredictable as suggested already in previous research, at larger scale [64,86,97–99], but also from an ethnographic perspective [96]. In such environments, hunter-gatherer groups would be less specialized in their hunting or foraging strategies, and carry a more versatile toolkit to cope with the unpredictable fluctuations in food resources. Thicker blades can be in fact used for a larger set of activities [60,85,86,88,89]. In light of these interpretations, we examine retouched tool assemblages as a complementary perspective on technological organization, comparing sites characterized by elongated, thinner blanks (more specialized) with those showing greater blank variability (less specialized and more flexible).

In GG1 from Enkapune Ya Muto, retouched tools prominently include large, backed segments, indicating a focus on robust tool forms, which are more abundant compared to those in the underlying assemblage. In DBL1, characterised by extremely high artefact densities, retouched tools are relatively rare, consisting of end-scrapers, scaled pieces, and comparatively to GG1, fewer geometric microliths, reflecting minimal retouching activity i.e. less maintenance and curation activities [50,54]. The Deka Wede 4 assemblage also features sparse retouched tools, including convergent sidescrapers, unifacial points, and end-scrapers. Some blanks display inverse proximal thinning retouch, but most artifacts were likely used without further modification, showing a tool-oriented production and a low degree of edge modification [19,49].

In the Middle and Late Holocene levels of Goda Buticha, retouched tools mainly consist of microliths, particularly backed microliths and retouched bladelets, predominantly made from obsidian, with a strong preference for specialized elongated blanks, these include unifacial and bifacial points with invasive retouch, as well as geometric microliths, such as crescents. This pattern is already visible in the late MSA levels of Goda Buticha (IId-IIf), retouched tools emphasize elongated blanks, particularly retouched blades made from obsidian. However, points, including unifacial and bifacial types, are significant and exhibit invasive retouch, showing a certain level of flexibility in the management of the tool-kit. At Gotera, only a small portion of the lithic assemblage is retouched, consisting mainly of notches and denticulates on blade and bladelet supports, primarily made of quartz [100,101]: this shows the production of a specialized tool-kit and a low degree of flexibility in the assemblage

Our analyses highlight the need to explore all aspects related to the interpretation of an archaeological context: not only the context and modality of the production but following all the stages of the use-life of the lithic objects, giving insight about mobility, duration of occupations, and object functions. Functional analysis could be a good complement to test the hypotheses emerging from these analyses, to better refine hunting strategies and connected hafting technologies [35,102]. Our results show the advantages of data integration and comparative analyses of lithic attributes to test general models built on the interpretation of individual sites to pursue this aim. There is an increasing necessity to standardize data collection for lithic industries and to establish a list of essential attributes to record for each assemblage. This standardization may facilitate the comparison not only for morphometric attributes but also technological traits, including reduction methods, techniques, etc.

Lastly, we still need to be cautious with the combination and use of archaeological data for large comparative analyses, especially when manipulating data on lithic technologies. We need a very deep understanding of these materials, the way they have been produced and the implications that technologies had for the reconstruction of past human behavior.

Our research represents the first case study in which the sites under consideration have been integrated into a large-scale approach based on the analysis of lithic assemblage attributes.

However, despite the results obtained, it is important to emphasize the need to incorporate more contexts within this model, following the same methodological approach, to achieve a large-scale understanding of the MSA-LSA transition in eastern Africa. In particular, it is necessary to mention several contexts that fall within this chronological range and would be essential to include in these analyses, to further test these patterns, but are not available at the time of this study. In Ethiopia sites dated to MIS 4 such as Shinfa Metema 1 [103]; to the MIS 3 such as Mochena Borago [104,105], Fincha Habera [106]; Gorgora [107]; to the MIS 2 such as Sodicho [108]. In Kenya sites dated to MIS 4 such as Panga ya Saidi [20,31], to the MIS 3 such as Rusinga Island sites [109]; Lukenya Hill [110]. This will allow to explore further results for the human technical behavior between MIS 5 and 1 in eastern Africa.

## Conclusions

Late Pleistocene in eastern Africa represents an ideal context to test different hypotheses linked to human group behavior, the reasons behind technological changes, and their relationship with climate and environment. The region is characterized by a diversity of environments that generated varied responses to past climatic fluctuations, leading to a differentiation in the behavioral responses of hunter-gatherers [4,17,41,111]. Our study shows a correlation between the shape of elongated blanks (and so, most likely, their use) with the harshness of the dry season. The available data shown here lead us to propose a model to further test in the future, i.e. that during periods in which the dry season is milder and the environment is more homogeneous through the year, longer and thinner blades and bladelets may be prevalent due to predictable resource availability. Because of such stability, hunting would have been more predictable, leading to a more specialized toolkit. In contrast, in moments with harsher and more challenging dry seasons, leading to a reduced resource availability for part of the year, shorter and thicker blades and bladelets may have been favored, reflecting a greater versatility in foraging strategies as well as other activities, to cope with unpredictable environments.

Through the combination of different data, both archaeological and paleoclimatic, with the collaboration of multiple researchers providing the raw data for the analyses, we have shown the importance of this study in explaining phenomena related to the cultural evolution of past human groups. Furthermore, our research highlighted the strong need to establish a fundamental list of attributes to record during analyses of lithic industries, enabling not only data sharing but also their comparison for minimizing errors. Additionally, it is desirable to strengthen the analyses on all aspects of archaeological data, including functional analyses of artifacts to gain a better understanding of the reasons behind some of the major technological changes in the archaeological record.

## Materials and methods

### Lithic assemblages

In this article, we have examined elongated unretouched blanks (named blades and bladelets according to the common terminology for lithic categorization, where they are defined as flakes with a length twice or more their width) from five archaeological sites in eastern Africa, spanning stratigraphic sequences from MIS 5 to MIS 1.

One reason for the lack of a clear model on factors driving technological changes in parts of eastern Africa is the limited amount of datable archaeological evidence from this period. On the other hand, there is also a complete lack of a comparative analysis of lithic assemblages that goes beyond the mere typological classification of artifacts, potentially leading to interpretative errors [112].

Based on the above statements, the sample selection for this analysis was based on two specific criteria:

1. **Chronological**; All the earliest MSA contexts (MIS 7-6) were excluded from the selection. Additionally, to integrate paleoclimatic data, a chronological attribution is needed.
2. **Analytical**. This is a fundamental criterion used to select samples for analysis. As mentioned in the introduction of this chapter, it is crucial to avoid typological comparisons based on the presence/absence of tool types that might exclude information about the production of lithic artifacts. To be included in a statistical methodology that compares technological attributes (both qualitative and quantitative), the collections and datasets must be amenable to a piece-by-piece artifact analysis in which all critical variables can be measured and assessed. It is clear that the technological approach based on the analysis of lithic artifact attributes is not a new methodology, however some assemblages collected and studied many years ago would require a full re-analysis to enable comparable data to be derived.

Based on the above criteria, in order to examine an archaeologically and chronologically significant sample, the sites of Goda Buticha, Porc Epic, Gotera 10, Deka Wede 4 in Ethiopia, and Enkapune ya Muto in Kenya were considered in this analysis.

### Archaeological sites

Goda Buticha is a cave in south-eastern Ethiopia with a long stratigraphic sequence divided into two main complexes, II and I. The deposit is dated from the end of MIS 4 to MIS 1, with a chronological gap dated to MIS 2 [19,113]. This site stands out as one of the few well-dated long stratigraphic sequences in Ethiopia [19].

Porc Epic cave, located in south-eastern Ethiopia, exhibits four main stratigraphic units. The lowest units (IV and III) are considered MSA based on the analysis of lithic assemblages; unit II shows a mixed MSA/LSA presence, while unit I is attributed to the LSA [49]. Here only artifacts from the lower units are considered in the analysis. Preliminary results of an OSL dating program of the lower units of Porc-Epic gave ages ranging from MIS 5 to MIS 4 [114,115]. We therefore worked with the hypothesis that the lower assemblages have an age range between 130 and 60 ka. We note that this chronological attribution is earlier than radiocarbon dates on shells available for the site[116,117] but other dates suggested an earlier chronology for the lower levels of Porc-Epic (see review in Assefa et al.[118]).

The Gotera area situated in southern Ethiopia near the Kenyan border has yielded open-air evidence of human occupation associated with MSA technological features [100,119]. The GOT-10 site has provided a stratigraphic sequence dated to MIS 3, including lithics, faunal remains, and evidence of fireplaces [101].

Deka Wede 4, located in the Ziway Sahla basin, is chronologically placed within MIS 3 through correlation with Deka Wede 1 site [120].

Lastly, Enkapune ya Muto, in southern Kenya, has yielded three levels dated to the Late Pleistocene: RBL4 containing MSA materials, GG and OL with LSA materials dating to ∼50 ka, and DBL dated around ∼40 ka with evidence of a younger LSA [50,54]. Here, only artifacts from GG and DBL layers were considered.

### Sources of raw materials

At Gotera, the principal raw materials used are quartz and basalt; they are available within the site, and in the surrounding area [100]. Obsidian sources in Ethiopia are relatively abundant [121]. Analyses conducted on obsidian artifacts from Porc Epic site showed that its source was located ∼250 km distant from the site. This evidence shows that this raw material was transported to the site from a very distant location [115]. In Goda Buticha Chert, basalt, and quartzite are assumed to have a local origin, while obsidian, despite its high frequency, lacks a known nearby source [19,122]. There is an obsidian outcrop c. 40 km away from the site, but its use has not been demonstrated [19]. Although in Goda Buticha there is no particular evidence for different raw material selection in the layer IIdIIf, in the IIc and I assemblages obsidian is the most used raw material, to produce both retouched tools and elongated blanks (only in GB-I) [19].

Lithic assemblages in Enkapune ya Muto show a few non-local raw materials but are mainly produced using obsidian from the volcanic area of the central rift, relatively close to the site [28].

## Data analysis

Regarding lithic categories (e.g., flakes, elongated flakes, tools, cores, etc.), we specifically chose to focus on unretouched and complete blades and bladelets. The reason is that their presence seems to increase during the LSA [29,49]. Moreover, they are the type of blanks for which the most data were available for the sites of interest, which allows us to increase the chronological and geographical scale of the analysis.

Each lithic in the dataset has been associated with a chronology and a cultural phase. Cultural phases have been attributed in accordance with the publication related to the contexts analyzed. Accordingly, we have identified the Middle Stone Age (MSA) for Porc Epic layers and the Goda Buticha layers dated between 63 and 31 ka; the Late Middle Stone Age (LMSA) for the Gotera 10 and Deka Wede 4 assemblages; the Late Stone Age (LSA) assemblages belong to Enkapune ya Muto show different chronologies which overlaps with some LMSA assemblages, in fact GG1 LSA layer at Enkapune ya Muto is dated ∼50 ka, while the DBL1 is dated ∼39ka. The Holocene assemblages (Middle and Late Holocene) from Goda Buticha, are dated from ∼8 ka to 700 years ago (Table 1) [19,49,51,52,100,113,119,120].

### Choice of technological attributes for the analyses

It is important to highlight that information on the lithic assemblages analyzed have been collected by different researchers (N=5) from various research traditions, each with distinct objectives. This led to great variability in the metrics considered, the categories analyzed, and other technical choices that can highly impact our analyses.

Caution must be given when choosing metrics and technological attributes to compare such different sets of data [56,112]. Following the approach designed by Pargeter and colleagues [56], based on a rigorous experiment demonstrating the statistical validity of certain attributes over others when the same assemblage is analyzed by different researchers. Therefore, we focused on specific metrics and technological attributes that are most suitable for statistical analyses of data collected by different individuals or research groups (Table 2). These include technological measurements (maximum length, width, and thickness), the number of dorsal scars, blank completeness, elongated index, and flattening index. We analyzed these attributes from the five different datasets related to the sites mentioned above.

**Table 2:** Variables used for the PCA on elongated blanks.

| Variables for elongated blanks |  |
| --- | --- |
| Maximum Length | XTL |
| Maximum Breadth | XTB |
| Maximum thickness | XT |
| Number of Scars | NR |
| Elongation index | EI |
| Flattening index | FI |

### Principal Component Analysis

We performed Principal Component Analysis on lithic attributes (Table 2) using the *FactoMinerR* package [123] and its *prcomp()* function. Cultural phases and stratigraphic units of each site were chosen as variables for grouping the PCA results and the two principal components. To better visualise how the different cultural phases are located in the PCA space, we computed the centroids for each archaeological layer (Figure 2) and raw material (figure 3) as the mean of PC1 and PC2 of all artifacts included in the respective category.

### Paleoclimate

To investigate the relationship between lithic attributes and the climatic conditions at each site at the time of occupation, we accessed and handled paleoclimatic data using the software *pastclim* [59], an R package that facilitates extracting and manipulating climatic data and paleoclimatic reconstructions. The package includes several sets of palaeoclimatic reconstructions ranging from a few thousand to millions of years (full list at https://evolecolgroup.github.io/pastclim/articles/a1_available_datasets.html). Given a set of geographic coordinates and their associated ages, *pastclim* retrieves reconstructed bioclimatic variables from the associated grid cell in the desired reconstruction.

For our analyses, we used the paleoclimatic simulations developed by Krapp et al. [58]. Those are a statistically-derived climatic dataset for the whole world covering the last 800,000 years at intervals of 1,000 years. As detailed in the original publication, it is based on a set of linear regressions between 72 existing HadCM3 climate simulations of the last 120,000 years and external forcings consisting of CO_2_, orbital parameters, and land type. The results compare well with both the original HadCM3 simulations and long-term proxy records [58]. The estimated climatologies were interpolated to 0.5° resolution and bias-corrected using the delta method [124].

The dataset includes annual estimates of 17 bioclimatic variables (*e.g.* minimum and maximum temperature, total precipitation, temperature seasonality). It also includes monthly temperature, precipitation, cloud cover, net primary productivity and global biome distributions at a resolution of 0.5° (degrees). It has been shown that in most cases downscaling such models to higher resolution has a minimal net effect on their consistency with pollen records [61], thus making it problematic to validate any high-resolution downscaling.

### Chronological resampling and paleoclimatic analyses

Some of the assemblages analyzed (Table 1) show a large chronological uncertainty. Although some sites among these are well dated—for example, Gotera, dated with AMS-¹⁴C [125], and Goda Buticha, dated using a combination of ¹⁴C and OSL methods [19,126], other sites, such as DW4, show an age based on correlation, and the dating of Porc-Epic remains debated. However, for Porc-Epic, we are using the new results from OSL dating [115]. The layers from Enkapune ya Muto considered here were dated using obsidian hydration and radiocarbon dating on ostrich eggshell [54].

Following Will et al 2021[65], for each assemblage we resampled randomly 1000 times a date from its whole chronological range associated.

We then used the R package *pastclim* [59] to access the paleoclimatic reconstructions from Krapp et al., 2021[58] (see section “Paleoclimate” above). For each location and resampled date, *pastclim* reports the variables of interest reconstructed for the associated 0.5×0.5 degrees grid cell, and 1000-years time slice.

We used the “*location_slice()*” function from *pastclim*, to associate each resampled set of dates for the assemblages with the relevant climate variables (i.e. reconstructed for their spatial coordinates and resampled age). We considered mean annual temperature (in the software coded as “bio01”), total annual precipitation (“bio12”), net primary productivity (“npp”) and precipitation of the driest season (“bio17”)

We then calculated 1000 times the Pearson’s correlation between the values of PC1 and PC2 for each centroid against the resampled time and associated climatic variables for their respective locations. We plotted each individual regression line, and calculated the mean value of rho, together with the number of resampled dates for which the p-value was below 0.05 [65]. The scatterplots were created using the *smplot2* package [127]. All graphical representations were generated using the *ggplot2* R package [128].

To check if the impact of seasonality is the result to large-scale fluctuations in precipitation

### Check for an effect of sample sizes over the results

The final dataset, created to be as comprehensive as possible, differs in terms of sample sizes (Table 3). The layer with the lowest number of analyzed artifacts (Goda Buticha IIdIIf 030-000) only includes 14 elongated blanks, while the layer with the biggest set of data is Goda Buticha IIc (160-100), with 186.

**Table 3:**
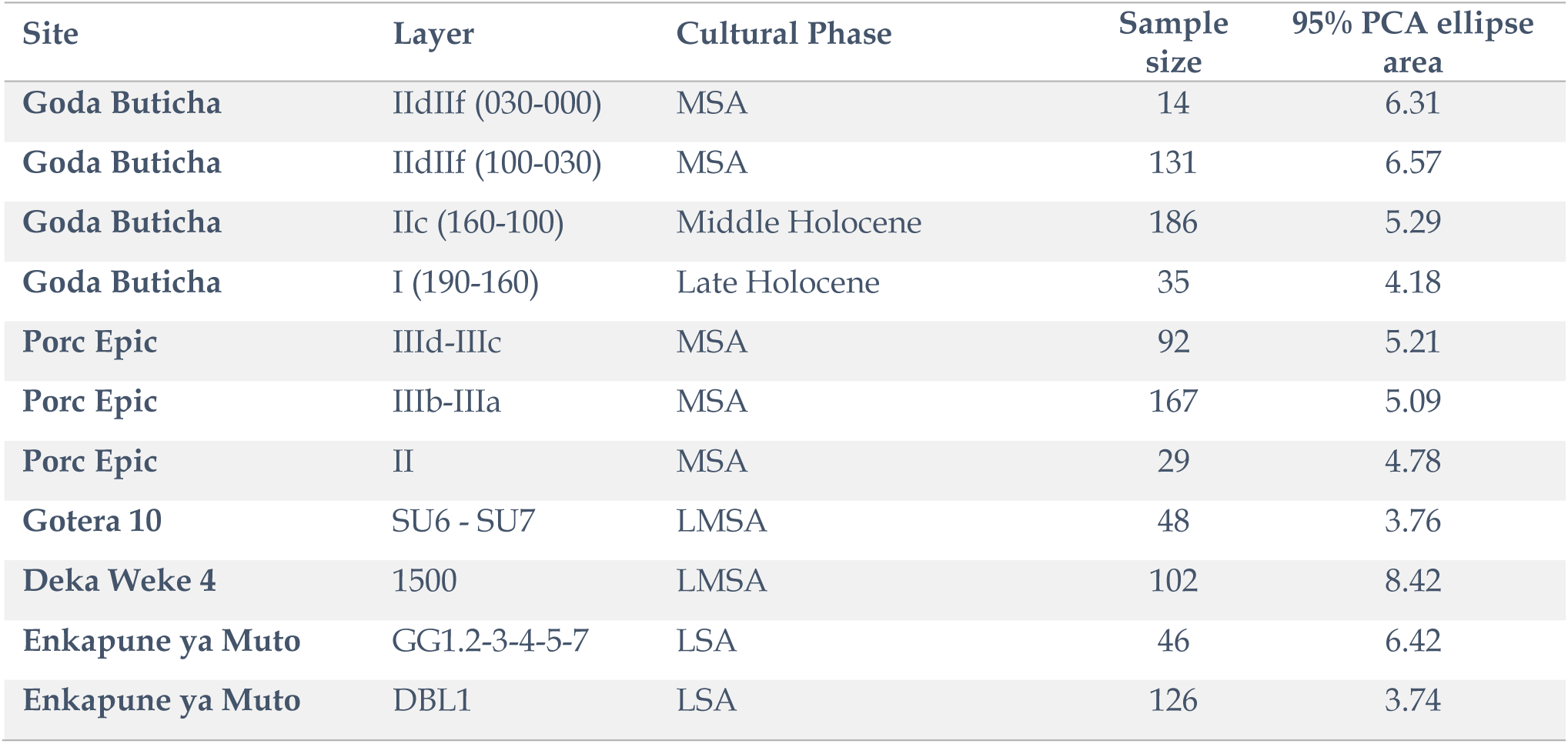
list of layers considered in the analysis, including the cultural phase, the number of elongated blanks, and the area of the associated ellipse including 95% of the points in the PCA space.

Variations in sample size may influence the results presented here; therefore, we conducted several tests to examine this possibility.

As an initial step, we assessed whether smaller sample sizes were associated with reduced diversity in the size and shape of the blanks. If so, these samples might have captured only a limited subset of the overall variability.

As an estimate of overall diversity in the lithic attributes within each assemblage, we calculated the area of the ellipse encompassing 95% of the data points in PCA space. As shown in Table 3 smaller sample sizes are not linked to reduced diversity. As an example, Goda Buticha IIdIIf 030-000 (sample size=14, the smallest in our dataset) is associated with an area of 6.31, which is larger to the one represented by the largest assemblage analysed, Goda Buticha IIc (160-100) (ellipse area= 5.26, sample size=186). A formal correlation test between sample size and ellipse area (Pearson’s ρ = 0.06, p-value = 0.73) provides no evidence of a significant association, indicating that smaller samples do not necessarily reflect lower diversity in the lithic attributes analyzed.

As a second step, we tested if sample sizes or ellipse areas show an association with the resampled ages (figure 6). Also in this case, correlations are low and not significant.

**Figure 6:**
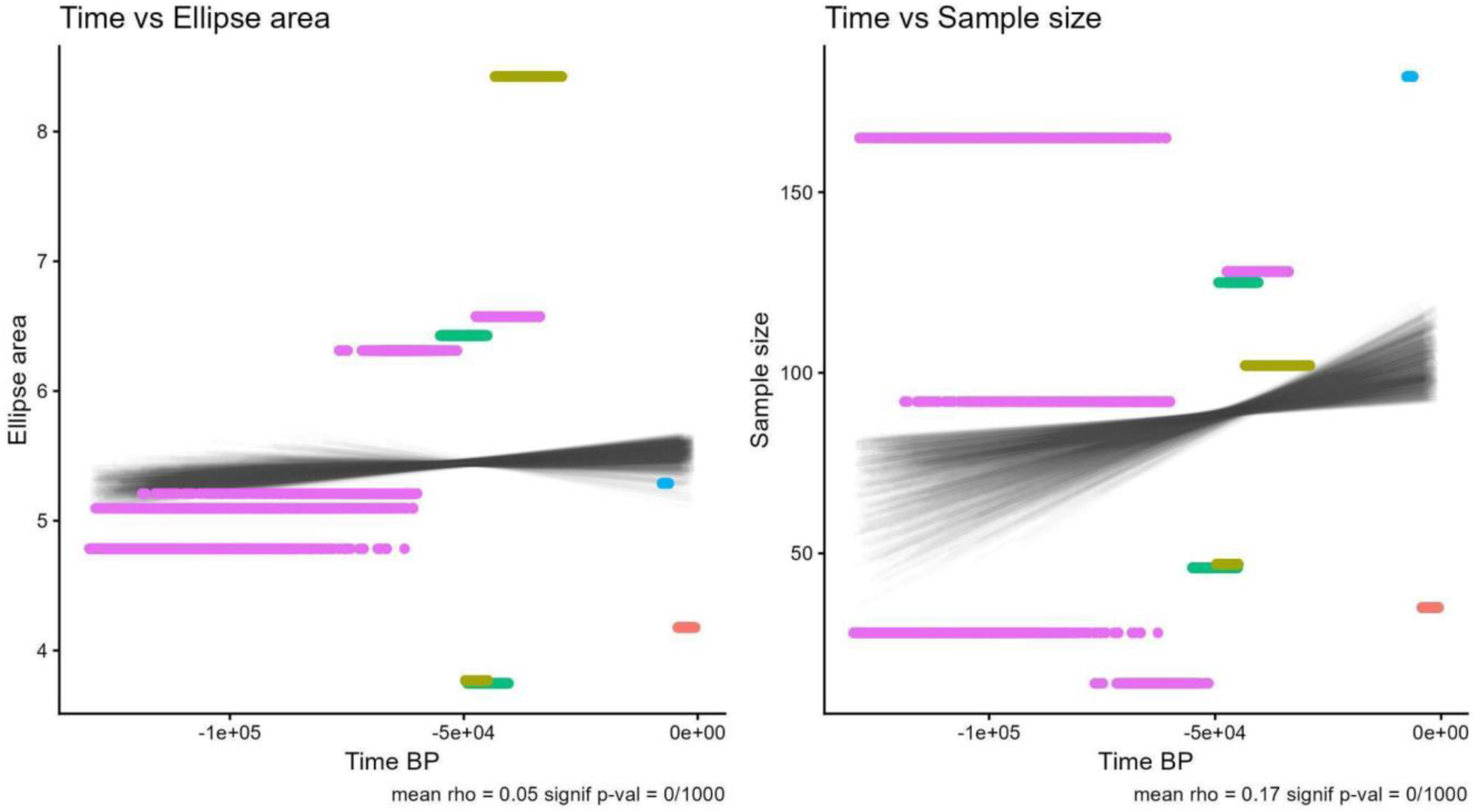
plot showing the non-significant correlation between the area of the ellipse including 95% of the points in the PCA space and time (left) and sample size and time (right).

As a result, we suggest that sample size does not appear to have a direct effect on the overall diversity of the assemblages, and is not apparently driven by time, supporting the validity of our analyses.

## Acknowledgements

We thank the Ethiopian Heritage Authority (EHA), the Borana Zone Culture and Tourism Office for their cooperation and permission to conduct fieldwork at Gotera. We thank EHA for permission to study lithic collections stored in Addis Ababa and the National Museums of Kenya in Nairobi for permission to study lithic collections from EYM.

We are grateful to Prof. Stan Ambrose, to Dr. Katja Douze for their help and contribution in studying lithics from EYM and DW4 respectively and to Dr. Clement Ménard for giving us access to DW4 lithic attributes.

We thank Francesco Lucchini for producing Figure 1.

## Author contributions

AM, ML and MF conceived the study with inputs from EES. MF, BH, AL collected the data. AM and ML defined the methodology. MF performed the analyses and created the figures with inputs from ML and AM. MF, ML, AM and EES provided the interpretation. MF and ML wrote the original manuscript. All authors were involved in reviewing the manuscript. MF, EES, ML and AM obtained the fundings.

## Data and code

Data and code needed to reproduce the analyses presented here have been uploaded to figshare: https://doi.org/10.6084/m9.figshare.28009085.v1. This includes the full dataset of lithic measurements, the associated climate variables, and all R scripts used for the analyses.

## References

1. Hovers, E. & Kuhn, S. Transitions before the Transition: Evolution and Stability in the Middle Paleolithic and Middle Stone Age. (2006).

2 . Leplongeon, A. et al. The Horn of Africa at the end of the Pleistocene (75 12 ka) in its macroregional context. in Not just a Corridor. Human occupation of the Nile Valley and neighbouring regions between 75,000 and 15,000 years ago (eds. Leplongeon, A., Goder-Goldberger, M. & Pleurdeau, D.) vol. Natures en Sociétés 269–283 (Muséum national d’Histoire naturelle, Paris, 2020).

3. Mcbrearty, S. & Brooks, A. S. The revolution that wasn’t: a new interpretation of the origin of modern human behavior. Journal Human Evolution 39, 453–563 (2000).

4. Tryon, C. A. The Middle/Later Stone Age transition and cultural dynamics of late Pleistocene East Africa. Evolutionary Anthropology vol. 28 267–282 Preprint at 10.1002/evan.21802 (2019).

5. Ambrose, S. H. Late Pleistocene Human Population Bottlenecks, Volcanic Winter, and Differentiation of Modern Humans. (1998).

6. Klein, R. G. Archeology and the Evolution of Human Behavior. *Evolutionary Anthropology: Issues*, News, and Reviews 17–36 (2000).

7. Mellars, P. Major Issues in the Emergence of Modern Humans. Curr. Anthropol. 30, 349–385 (1989).

8. Mellars, P. The impossible coincidence. A single-species model for the origins of modern human behavior in Europe. Evolutionary Anthropology vol. 14 12–27 Preprint at 10.1002/evan.20037 (2005).

9 . Mellars, P. The Neanderthal Legacy: An Archaeological Perspective from Western Europe. (Princeton University Press., 1998).

10. Jones, R., Mellars, P. & Stringer, C. The Human Revolution: Behavioural and Biological Perspectives on the Origins of Modern Humans. (Princetown University Press, 1989).

11. White, M. & Ashton, N. Lower Palaeolithic core technology and the origins of the Levallois method in North-Western Europe. Curr. Anthropol. 44, 598–608 (2003).

12. Scerri, E. M. L. & Will, M. The revolution that still isn’t: The origins of behavioral complexity in Homo sapiens. J. Hum. Evol. 179, 103358 (2023).

13. d’Errico, F. et al. Trajectories of cultural innovation from the Middle to Later Stone Age in Eastern Africa: Personal ornaments, bone artifacts, and ocher from Panga ya Saidi, Kenya. J. Hum. Evol. 141, 102737 (2020).

14. Martinón-Torres, M. et al. Earliest known human burial in Africa. Nature 593, 95–100 (2021).

15. Wilkins, J. et al. Innovative Homo sapiens behaviours 105,000 years ago in a wetter Kalahari. Nature 592, 248–252 (2021).

16. Brooks, A. S. et al. Long-distance stone transport and pigment use in the earliest Middle Stone Age. Science (1979). 360, 90–94 (2018).

17. Leplongeon, A. et al. A comparative look at technical traditions in the Horn of Africa and the Nile Valley at the end of the Pleistocene. in Actes de la Séance de la Société Préhistorique Française de Toulouse les 25-28 septembre 2019 (Société Préhistorique française, Paris, 2019).

18. Brandt, S. A. The Upper Pleistocene and early Holocene prehistory of the Horn of Africa. Afr. Archaeol. Rev. 4, 41–82 (1986).

19. Leplongeon, A., Pleurdeau, D. & Hovers, E. Late pleistocene and holocene lithic variability at Goda Buticha (Southeastern Ethiopia): Implications for the understanding of the middle and late stone age of the horn of Africa. Journal of African Archaeology 15, 202–233 (2017).

20. Shipton, C. et al. 78,000-year-old record of Middle and Later stone age innovation in an East African tropical forest. Nat. Commun. 9, (2018).

21. Goodwin, Astley John Hilary; Van Riet Lowe, C. The Stone Age Cultures of South Africa. (1929).

22. Hublin, J. J. et al. New fossils from Jebel Irhoud, Morocco and the pan-African origin of Homo sapiens. Nature 546, 289–292 (2017).

23. Richter, D. et al. The age of the hominin fossils from Jebel Irhoud, Morocco, and the origins of the Middle Stone Age. Nature 546, 293–296 (2017).

24. Fernández, V. M., de la Torre, I., Luque, L., González-Ruibal, A. & López-Sáez, J. A. A Late Stone Age sequence from West Ethiopia: the sites of K’aaba and Bel K’urk’uma (Assosa, Benishangul-Gumuz Regional State). Journal of African Archaeology 5, 91–126 (2007).

25. Gutherz, X. et al. The Hargeisan revisited: Lithic industries from shelter 7 of Laas Geel, Somaliland and the transition between the Middle and Late Stone Age in the Horn of Africa. Quaternary International 343, 69–84 (2014).

26. Scerri, E. M. L., Blinkhorn, J., Niang, K., Bateman, M. D. & Groucutt, H. S. Persistence of Middle Stone Age technology to the Pleistocene/Holocene transition supports a complex hominin evolutionary scenario in West Africa. J. Archaeol. Sci. Rep. 11, 639–646 (2017).

27. Scerri, E. M. L. et al. Continuity of the Middle Stone Age into the Holocene. Sci. Rep. 11, 70 (2021).

28. Ambrose, S. H. Small Things Remembered: Origins of Early Microlithic Industries in Sub-Saharan Africa. Archaeological Papers of the American Anthropological Association 12, 9–29 (2002).

29. Grove, M. & Blinkhorn, J. Neural networks differentiate between Middle and Later Stone Age lithic assemblages in eastern Africa. PLoS One 15, (2020).

30. Grove, M. & Blinkhorn, J. Testing the Integrity of the Middle and Later Stone Age Cultural Taxonomic Division in Eastern Africa. Journal of Paleolithic Archaeology 4, 14 (2021).

31. Shipton, C. et al. The Middle to Later Stone Age transition at Panga ya Saidi, in the tropical coastal forest of eastern Africa. J. Hum. Evol. 153, 102954 (2021).

32. Tryon, C. A. & Faith, J. T. A demographic perspective on the middle to later stone age transition from nasera rockshelter, Tanzania. Philosophical Transactions of the Royal Society B: Biological Sciences 371, (2016).

33. Blinkhorn, J. & Grove, M. Explanations of variability in Middle Stone Age stone tool assemblage composition and raw material use in Eastern Africa. Archaeol. Anthropol. Sci. 13, 14 (2021).

34. Shipton, C. Miniaturization and Abstraction in the Later Stone Age. Biol. Theory 18, 253–268 (2023).

35. Basell, L. S. & Spinapolice, E. E. Time, the Middle Stone Age and lithic analyses following the Third Science Revolution. Azania: Archaeological Research in Africa 59, 140–159 (2024).

36. Hussain, S. T. & Soressi, M. The Technological Condition of Human Evolution: Lithic Studies as Basic Science. Journal of Paleolithic Archaeology 4, 25 (2021).

37. Basell, L. S. Middle Stone Age (MSA) site distributions in eastern Africa and their relationship to Quaternary environmental change, refugia and the evolution of Homo sapiens. Quat. Sci. Rev. 27, 2484–2498 (2008).

38. Beyer, R. M., Krapp, M., Eriksson, A. & Manica, A. Climatic windows for human migration out of Africa in the past 300,000 years. Nat. Commun. 12, 4889 (2021).

39. Blinkhorn, J., Timbrell, L., Grove, M. & Scerri, E. M. L. Evaluating refugia in recent human evolution in Africa. Philosophical Transactions of the Royal Society B: Biological Sciences 377, (2022).

40. Clark, J. D. The Middle Stone Age of East Africa and the beginnings of regional identity. J. World Prehist. 2, 235–305 (1988).

41. Blome, M. W., Cohen, A. S., Tryon, C. A., Brooks, A. S. & Russell, J. The environmental context for the origins of modern human diversity: A synthesis of regional variability in African climate 150,000-30,000 years ago. J. Hum. Evol. 62, 563–592 (2012).

42. Gasse, F., Chalié, F., Vincens, A., Williams, M. A. J. & Williamson, D. Climatic patterns in equatorial and southern Africa from 30,000 to 10,000 years ago reconstructed from terrestrial and near-shore proxy data. Quat. Sci. Rev. 27, 2316–2340 (2008).

43. Gasse, F. Hydrological changes in the African tropics since the Last Glacial Maximum. Quat. Sci. Rev. 19, 189–211 (2000).

44. Fischer, M. L. et al. Determining the Pace and Magnitude of Lake Level Changes in Southern Ethiopia Over the Last 20,000 Years Using Lake Balance Modeling and SEBAL. Front. Earth Sci. (Lausanne). 8, (2020).

45. Umer, M. et al. Late Quaternary climate changes in the Horn of Africa. in 159–180 (2004). doi:10.1007/978-1-4020-2121-3_9.

46. Marshall, M. H. et al. Late Pleistocene and Holocene drought events at Lake Tana, the source of the Blue Nile. Glob. Planet. Change 78, 147–161 (2011).

47. Scerri, E. M. L., Drake, N. A., Jennings, R. & Groucutt, H. S. Earliest evidence for the structure of Homo sapiens populations in Africa. Quat. Sci. Rev. 101, 207–216 (2014).

48. Timbrell, L. et al. Stone point variability reveals spatial, chronological and environmental structuring of eastern African Middle Stone Age populations. Azania 59, 111–139 (2024).

49. Leplongeon, A. Microliths in the Middle and Later Stone Age of eastern Africa: New data from Porc-Epic and Goda Buticha cave sites, Ethiopia. Quaternary International 343, 100–116 (2014).

50. Leplongeon, A. Middle Stone Age and early Late Stone Age lithic assemblages at Enkapune Ya Muto (Kenya). in Proceedings of the European Society for the study of Human Evolution. 141 (2016).

51. Ménard, C. et al. Late Stone Age variability in the Main Ethiopian Rift: New data from the Bulbula River, Ziway-Shala basin. Quaternary International 343, 53–68 (2014).

52. Ménard, C. & Bon, F. The Bulbula River Sites, Ethiopia. in Handbook of Pleistocene Archaeology of Africa 275–283 (Springer International Publishing, Cham, 2023). doi:10.1007/978-3-031-20290-2_16.

53. Khalidi, L. et al. 9000 years of human lakeside adaptation in the Ethiopian Afar: Fisher-foragers and the first pastoralists in the Lake Abhe basin during the African Humid Period. Quat. Sci. Rev. 243, (2020).

54. Ambrose, S. H. Chronology of the later stone age and food production in East Africa. J. Archaeol. Sci. 25, 377–392 (1998).

55. Diez-Martín, F. et al. The Middle to Later Stone Age Technological Transition in East Africa. New Data from Mumba Rockshelter Bed V (Tanzania) and Their Implications for the Origin of Modern Human Behavior. Source: Journal of African Archaeology vol. 7 (2009).

56. Pargeter, J. et al. Replicability in Lithic Analysis. Am. Antiq. 88, 163–186 (2023).

57. Pargeter, J. & Brandt, S. A. Lithic Technological Approaches to the African Late Pleistocene Later Stone Age. Evolutionary Anthropology vol. 24 167–169 Preprint at 10.1002/EVAN.21461 (2015).

58. Krapp, M., Beyer, R. M., Edmundson, S. L., Valdes, P. J. & Manica, A. A statistics-based reconstruction of high-resolution global terrestrial climate for the last 800,000 years. Sci. Data 8, 228 (2021).

59. Leonardi, M., Hallett, E. Y., Beyer, R., Krapp, M. & Manica, A. *pastclim* 1.2: an R package to easily access and use paleoclimatic reconstructions. Ecography 2023, (2023).

60. Bleed, P. The optimal design of hunting weapons: maintainability or reliability. Am. Antiq. 51, 737–747 (1986).

61. Timbrell, L. et al. More is not always better: delta-downscaling climate model outputs from 30 to 5 min resolution has minimal impact on coherence with Late Quaternary proxies. Climate of the Past 21, 1185–1208 (2025).

62. Gosling, W. D. et al. A multi-model approach to the spatial and temporal characterization of the African Humid Period. Quaternary International 744, 109933 (2025).

63. Thomson, D. F. The seasonal factor in human culture illustrated from the life of a contemporary nomadic group. Proceedings of the Prehistoric Society 5, 209–221 (1939).

64. Timbrell, L. et al. Climate seasonality and predictability during the middle stone age and implications for technological diversification in early Homo sapiens. Sci. Rep. 15, (2025).

65. Will, M., Krapp, M., Stock, J. T. & Manica, A. Different environmental variables predict body and brain size evolution in Homo. Nat. Commun. 12, 4116 (2021).

66. Pargeter, J. & Shea, J. J. Going big versus going small: Lithic miniaturization in hominin lithic technology. *Evolutionary Anthropology: Issues*, News, and Reviews 28, 72–85 (2019).

67. Pargeter, J. Lithic miniaturization in Late Pleistocene southern Africa. J. Archaeol. Sci. Rep. 10, 221–236 (2016).

68. Kuhn, S. L. & Elston, R. G. Introduction: Thinking Small Globally. Archaeological Papers of the American Anthropological Association 12, 1–7 (2002).

69. Shea, J. J. The Middle Stone Age archaeology of the Lower Omo Valley Kibish Formation: Excavations, lithic assemblages, and inferred patterns of early Homo sapiens behavior. J. Hum. Evol. 55, 448–485 (2008).

70. Yellen, J. E. et al. The archaeology of Aduma Middle Stone Age sites in the Awash Valley, Ethiopia. PaleoAnthropology 2005, 25–100 (2005).

71. Porraz, G. et al. Technological successions in the Middle Stone Age sequence of Diepkloof Rock Shelter, Western Cape, South Africa. J. Archaeol. Sci. 40, 3376–3400 (2013).

72. Soriano, S., Villa, P. & Wadley, L. Blade technology and tool forms in the Middle Stone Age of South Africa: the Howiesons Poort and post-Howiesons Poort at Rose Cottage Cave. J. Archaeol. Sci. 34, 681–703 (2007).

73. Wurz, S. Technological Trends in the Middle Stone Age of South Africa between MIS 7 and MIS 3. Curr. Anthropol. 54, S305–S319 (2013).

74. Hiscock, P., Clarkson, C. & Mackay, A. Big debates over little tools: ongoing disputes over microliths on three continents. World Archaeol. 43, 653–664 (2011).

75. Clark, J. D. Raw material and African lithic technology. Man and Environment 4, 44–55 (1980).

76. Eren, M. I., Roos, C. I., Story, B. A., von Cramon-Taubadel, N. & Lycett, S. J. The role of raw material differences in stone tool shape variation: An experimental assessment. J. Archaeol. Sci. 49, 472–487 (2014).

77. Wilson, L. & Browne, C. L. Change in raw material selection and subsistence behaviour through time at a Middle Palaeolithic site in southern France. J. Hum. Evol. 75, 28–39 (2014).

78. Pargeter, J. & Faith, J. T. Lithic miniaturization as adaptive strategy: a case study from Boomplaas Cave, South Africa. Archaeol. Anthropol. Sci. 12, 225 (2020).

79. Scholes, R. J., & Archer, S. R. Tree–grass interactions in savannas. Annual Review of Ecology and Systematics, 28, 517–544 (1997).

80. Ogutu, J. O., Piepho, H. P., Dublin, H. T., Bhola, N., & Reid, R. S. Dynamics of Mara–Serengeti ungulates in relation to rainfall. Ecological Monographs, 78(2), 233–254 (2008).

81. Clark, J. & Linares-Matás, G. J. Seasonality and Lithic Investment in the Oldowan. Journal of Paleolithic Archaeology 6, (2023).

82. Linares-Matás, G. J. & Clark, J. Seasonality and Oldowan behavioral variability in East Africa. J. Hum. Evol. 164, 103070 (2022).

83. Jones, E. L. Mobility, settlement, and resource patchiness in Upper Paleolithic Iberia. Quaternary International 318, 46–52 (2013).

84. Grove, M. Hunter–gatherer movement patterns: Causes and constraints. J. Anthropol. Archaeol. 28, 222–233 (2009).

85. Nelson, M. C. The Study of Technological Organization. Source: Archaeological Method and Theory vol. 3 https://about.jstor.org/terms (1991).

86. Clark, J. & Linares-Matás, G. J. When to generalise and when to specialise? Climate change and hominin biocultural adaptability in the African early and middle stone age. Quaternary Science Advances 15, (2024).

87. Bamforth, D. B. Technological efficiency and tool curation. Am. Antiq. 51, 38–50 (1986).

88. Binford, L. R. Organization and Formation Processes: Looking at Curated Technologies. Source: Journal of Anthropological Research vol. 35 (1979).

89. Binford, L. R. Willow Smoke and Dogs’ Tails: Hunter-Gatherer Settlement Systems and Archaeological. Source: American Antiquity vol. 45 https://www.jstor.org/stable/279653 (1980).

90. Parry, W. J. Expedient core technology and sedentism. in The organization of core technology (eds. Johnson, J. K. & Morrow, C. A.) 285–304 (1987).

91. Shott, M. J. On Tool-Class Use Lives and the Formation of Archaeological Assemblages. Source: American Antiquity vol. 54 https://about.jstor.org/terms (1989).

92. Shott, M. J. An Exegesis of the Curation Concept. Source: Journal of Anthropological Research vol. 52 https://www.jstor.org/stable/3630085 (1996).

93. Ebert, J. I. Alternative Strategies for the Transport, Curation, and Use of Lithic Materials and Implements:“Discontinuous” Assemblages and Their Analytic Implications. in 46th Annual Meeting of the Society for American Archaeology, San Diego. (1979).

94. Kelly, R. L. Hunter-Gatherer Mobility *Strategies*. Source: Journal of Anthropological Research vol. 39 https://www.jstor.org/stable/3629672 (1983).

95. Torrence, R. Time budgeting and hunter-gatherer technology. Hunter-gatherer economy in prehistory 11–22 (1983).

96. Linares Matás, G. J. & Lim, J. S. “This is the way”: Knowledge networks and toolkit specialization in the circumpolar coastal landscapes of western Alaska and Tierra del Fuego. Journal of Island and Coastal Archaeology 19, 1–29 (2024).

97. Grove, M. Change and variability in Plio-Pleistocene climates: modelling the hominin response. J. Archaeol. Sci. 38, 3038–3047 (2011).

98. Potts, R. Evolution and Climate Variability. Science (1979). 273, (1996).

99 . Maslin, M. A. et al. East african climate pulses and early human evolution. Quaternary Science Reviews vol. 101 1–17 Preprint at 10.1016/j.quascirev.2014.06.012 (2014).

100. Fusco, M., Carletti, E., Gallinaro, M., Zerboni, A. & Spinapolice, E. E. Lithic variability and raw material exploitation at the Middle Stone Age (MSA) site of Gotera, southern Ethiopia: A combined technological and quantitative approach. Journal of Lithic Studies 8, (2021).

101. Fusco, M. et al. The Gotera Archaeological Mission in Southern Ethiopia: A preliminary field report on the ongoing research at the Middle Stone Age site of Gotera. Annales d’Ethiopie 35, (2024).

102. Rots, V. & Coppe, J. Functional perspectives on the Evolution of Hunting Technology in Africa and Europe. in Annual meeting of the Society for American Archaeology (2024).

103. Kappelman, J. et al. Adaptive foraging behaviours in the Horn of Africa during Toba supereruption. Nature 628, 365–372 (2024).

104. Brandt, S., Hildebrand, E., Vogelsang, R., Wolfhagen, J. & Wang, H. A new MIS 3 radiocarbon chronology for Mochena Borago Rockshelter, SW Ethiopia: Implications for the interpretation of Late Pleistocene chronostratigraphy and human behavior. J. Archaeol. Sci. Rep. 11, 352–369 (2017).

105. Brandt, S. A. et al. Early MIS 3 occupation of Mochena Borago Rockshelter, Southwest Ethiopian Highlands: Implications for Late Pleistocene archaeology, paleoenvironments and modern human dispersals. Quaternary International 274, 38–54 (2012).

106. Ossendorf, G. et al. Middle Stone Age foragers resided in high elevations of the glaciated Bale Mountains, Ethiopia. Science (1979). 365, 583–587 (2019).

107. Sahle, Y., Firew, G. A., Pearson, O. M., Stynder, D. D. & Beyin, A. MIS 3 innovative behavior and highland occupation during a stable wet episode in the Lake Tana paleoclimate record, Ethiopia. Sci. Rep. 14, 17038 (2024).

108. Bodin, S. C. et al. Afromontane forests and human impact after the African Humid Period: wood charcoal from the Sodicho rock shelter, SW Ethiopian highlands. Veg. Hist. Archaeobot. 33, 529–543 (2024).

109. Tryon, C. A. Lake Victoria Basin, Kenya. in Handbook of Pleistocene Archaeology of Africa 607–621 (Springer International Publishing, Cham, 2023). doi:10.1007/978-3-031-20290-2_38.

110. Tryon, C. A. et al. Late Pleistocene age and archaeological context for the hominin calvaria from GvJm-22 (Lukenya Hill, Kenya). Proceedings of the National Academy of Sciences 112, 2682–2687 (2015).

111. Timbrell, L., Grove, M., Manica, A., Rucina, S. & Blinkhorn, J. A spatiotemporally explicit paleoenvironmental framework for the Middle Stone Age of eastern Africa. Sci. Rep. 12, 3689 (2022).

112. Will, M. et al. Comparative analysis of Middle Stone Age artifacts in Africa (CoMSAfrica). Evol. Anthropol. 28, 57–59 (2019).

113. Pleurdeau, D. Human technical behavior in the African middle stone age: the lithic assemblage of Porc-Epic cave (Dire Dawa, Ethiopia). African Archaeological Review 22, 177–197 (2005).

114. Leplongeon, A., Goder-Goldberger, M., Pleurdeau, D. & Tribolo, C. Porc-Epic Cave, more than 90 years later – a typical MSA site in the Horn of Africa? in 16th Congress of the Pan African Archaeological Association for Prehistory and related studies (Zanzibar, 2022).

115. Tribolo, C., Leplongeon, A., Pleurdeau, D. & Assefa, Z. Applying isochrones methods with the post IR-IR290 signal for dating Middle Stone Age sites. in German Luminescence dating conference (DLED 2022) (Bonn, Germany, 2022).

116. Assefa, Z., Lam, Y. M. & Mienis, H. K. Symbolic use of terrestrial gastropod opercula during the Middle Stone Age at Pore-Epic cave, Ethiopia. Curr. Anthropol. 49, 746–756 (2008).

117. Rosso, D. E., D’Errico, F. & Queffelec, A. Patterns of change and continuity in ochre use during the late Middle Stone Age of the Horn of Africa: The Porc-Epic Cave record. PLoS One 12, (2017).

118. Assefa, Z., Pleurdeau, D., Leplongeon, A. & Lam, Y.-M. Porc-Epic Cave, Ethiopia. in Handbook of Pleistocene Archaeology of Africa 503–517 (Springer International Publishing, Cham, 2023). doi:10.1007/978-3-031-20290-2_31.

119. E.E. Spinapolice, M. Gallinaro, A. Z. New investigations in southern Ethiopia (Yabelo and Gotera): Pleistocene and Holocene archaeological evidences. Scienze dell’antichità 23, (2017).

120. Ménard, C. & Bon, F. The Bulbula River Sites, Ethiopia. in Handbook of Pleistocene Archaeology of Africa 275–283 (Springer International Publishing, Cham, 2023). doi:10.1007/978-3-031-20290-2_16.

121. Negash, A. & Shackley, M. S. Geochemical provenance of obsidian artefacts from the MSA site of Porc Epic, Ethiopia. Archaeometry 48, 1–12 (2006).

122. Pleurdeau, D. et al. Cultural change or continuity in the late MSA/Early LSA of southeastern Ethiopia? The site of Goda Buticha, Dire Dawa area. Quaternary International 343, 117–135 (2014).

123. Lê, S., Josse, J., Rennes, A. & Husson, F. FactoMineR: An R Package for Multivariate Analysis. JSS Journal of Statistical Software vol. 25 http://www.jstatsoft.org/ (2008).

124. Beyer, R. M., Krapp, M. & Manica, A. High-resolution terrestrial climate, bioclimate and vegetation for the last 120,000 years. Sci. Data 7, (2020).

125. Fusco, M. et al. First evidence of open-air human occupation dated to MIS3 in southern Ethiopia: the analysis of GOT-10 site, Gotera area. Quat. Sci. Rev.

126. Pleurdeau, D. et al. Goda Buticha, Ethiopia. in Handbook of Pleistocene Archaeology of Africa 337–352 (Springer International Publishing, Cham, 2023). doi:10.1007/978-3-031-20290-2_20.

127. Min, S. H. & Zhou, J. smplot: An R Package for Easy and Elegant Data Visualization. Front. Genet. 12, (2021).

128. Wickham, H. Ggplot2: Elegant Graphics for Data Analysis. (Springer-Verlag New York., 2016).

